# Nonadaptive Radiation: Pervasive diet specialization by drift in scale insects?

**DOI:** 10.1101/064220

**Authors:** Nate B Hardy, Daniel A Peterson, Benjamin B Normark

## Abstract

At least half of metazoan species are herbivorous insects. Why are they so diverse? Most herbivorous insects feed on few plant species, and adaptive host specialization is often invoked to explain their diversification. Nevertheless, it is possible that the narrow host ranges of many herbivorous insects are non-adaptive. Here, we test predictions of this hypothesis with comparative phylogenetic analyses of scale insects, a group for which there appears to be few host-use tradeoffs that would select against polyphagy, and for which passive wind-dispersal should make host specificity costly. We infer a strong positive relationship between host range and diversification rate, and a marked asymmetry in cladogenetic changes in diet breadth. These results are consonant with a system of pervasive non-adaptive host specialization in which small, drift-and extinction-prone populations are frequently isolated from persistent and polyphagous source populations. They also contrast with the negative relationship between diet breadth and taxonomic diversification that has been estimated in butterflies, a disparity which likely stems from differences in the average costs and benefits of host specificity and generalism in scale insects vs. butterflies. Our results indicate the potential for non-adaptive processes to be important to diet-breadth evolution and taxonomic diversification across herbivorous insects.

## Introduction

Species richness is spread unevenly across the branches of the Tree of Life. What causes this unevenness? One factor might be what a species eats. Herbivory has evolved repeatedly among insects, and herbivorous insects tend to be more species-rich than their non-herbivorous relatives (Mitter et al. 1988; Wiens et al. 2015; but see Rainford and Mayhew 2015). In fact, herbivorous insects are more species-rich than any other group of metazoans (Futuyma and Agrawal 2009). Understanding what drives species diversification in herbivorous insects is tantamount to understanding most of species diversification. Host-plant associations are thought to be key (Ehrlich and Raven 1964; Futuyma and Moreno 1988; Futuyma and Agrawal 2009).

Herbivorous insects vary conspicuously in their diet breadth. Most are host-plant specialists, but frequency distributions of host ranges are right-skewed (e.g., Fig 1), and herbivorous insect communities invariably include a minority of broad-diet (polyphagous) species (Forister et al. 2015). For the most part, we have assumed that host specialization is adaptive for herbivorous insects and we have incorporated this assumption into our explanations of herbivorous insect diversification (Ehrlich and Raven 1964). The Oscillation Hypothesis (Janz et al. 2006) is a good example. It explains herbivorous insect diversification as adaptive radiation (Yoder et al. 2010): an ecological opportunity decreases competition, relaxes stabilizing selection on host use, and causes the host range of a herbivorous insect species to expand. But host-range expansion is only temporary, because as competition rebounds, host-use trade-offs select against a broad diet and drive speciation through host specialization. Note that, in adaptive models, among-host performance trade-offs are essential; without them, there is no cost for polyphagy. But evidence for these trade-offs has been hard to find. In fact, most studies have found instead that within herbivorous insect populations “general vigor” genotypes outperform others across all viable hosts (e.g., Forister et al. 2007; Agosta and Klemens 2009; Gompert et al. 2015). Host-use tradeoffs are also called into question by the results of comparative phylogenetic analyses: in butterflies, host-use trade-offs do not select strongly enough against generalists so as to make broad diets evolutionarily ephemeral (Hardy and Otto 2014; Hamm and Fordyce 2015), and in armored scale insects host-use trade-offs do not constrain the phylogenetic history of host use (Peterson et al. 2015). This raises the possibility that much of the host specificity observed in herbivorous insects is not adaptive.

**Figure 1.**
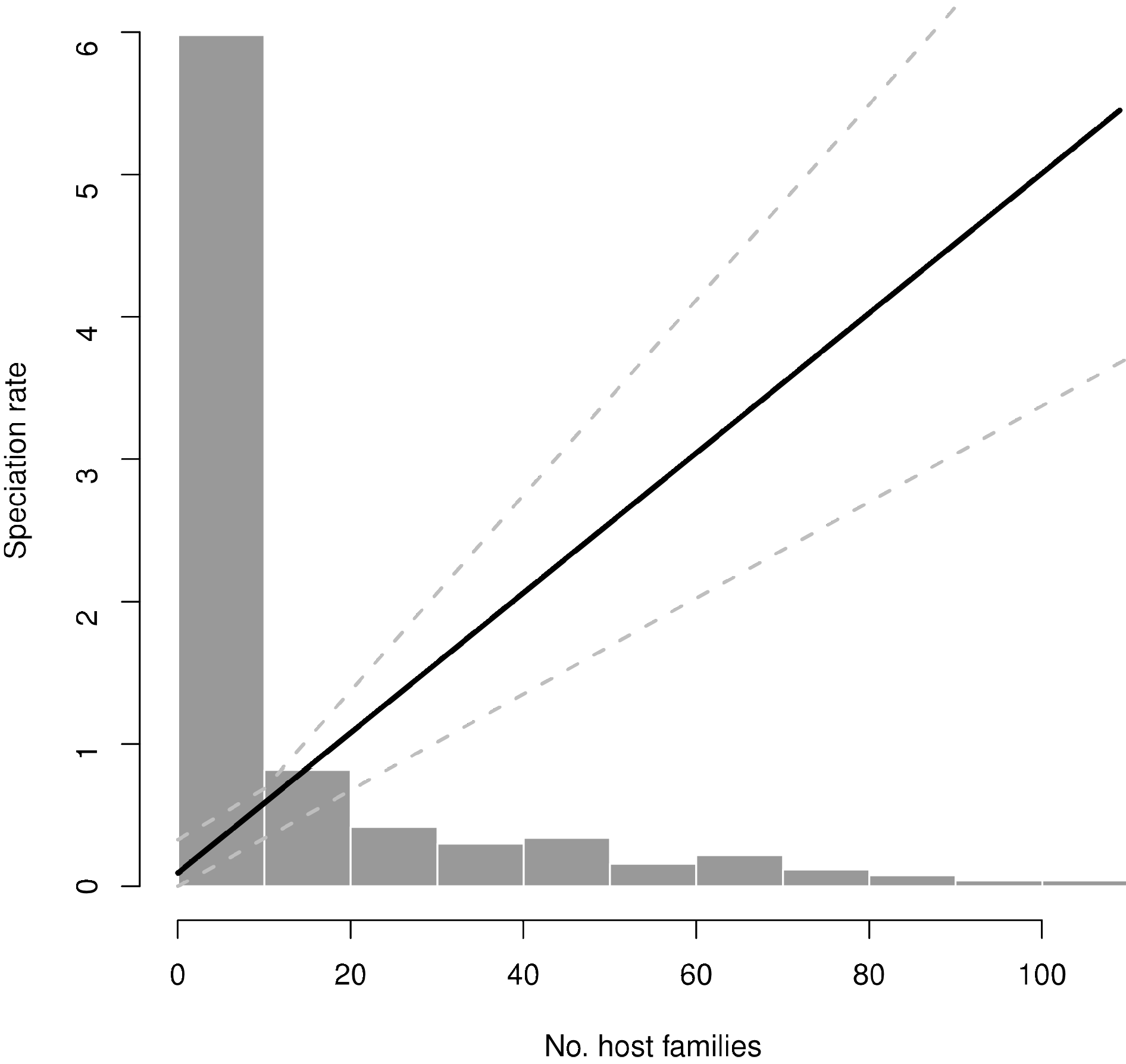
Speciation rate estimated as a function of host plant range (number of host families). The number of host families used by each of the 426 scale insect species examined is shown as a histogram. The solid black line shows the Bayesian estimate of the mean relationship between host range and speciation under a QuaSSE model (speciation rate = 0.095 + 0.050 * host range). The dashed gray lines show the 95% High Posterior Density.

Instead, host specificity may evolve by genetic drift (Gompert et al. 2015). Small populations may become geographically cut off from polyphagous source populations. In these isolated groups, selection for adaptive polyphagy may not be strong enough to overcome the power of genetic drift, or selection for polyphagy may be relaxed if ancestral hosts are absent in restricted geographic ranges. We refer to this possibility as the specialization-by-drift hypothesis. Regardless of how specialization comes about, it should increase the likelihood of extinction (Simpson 1953; Jablonski 1986). And if it evolves through genetic drift, it may be especially likely to reduce the potential for subsequent adaptive evolution, such as host switching or host range expansion (Jaenike 1990). Hence, the specialists produced by genetic drift can be thought of as evolutionary dead-ends. The idea of specialists as dead-ends is an old and conventional one in evolutionary biology (Cope 1896; Huxley 1942; Simpson 1953; Day et al. 2016); conventionally, specialists are thought to have a short-term advantage and long-term disadvantage. In the specialization-by-drift model they enjoy no advantage at all.

In short, the standard adaptive-specialization hypothesis views generalists as unstable and ephemeral, rather like radioactive isotopes that rapidly decay, whereas the specialization-by-drift hypothesis views generalists as stable and persistent, like a kind of phylogenetic meristem. Remarkably, although the reasons differ, the standard adaptive-specialization hypothesis and the specialization-by-drift hypothesis make many of the same predictions about the evolution of host range and its effects on species diversification. Both hypotheses predict that 1) host range evolution will be tightly linked to speciation events, 2) host ranges will shrink more than expand, and 3) generalists will speciate more rapidly than specialists.

To reiterate, in the standard version of the adaptive-specialization hypothesis, a rare ecological opportunity causes the host range of a herbivorous insect species to expand. But before long, escalating competition renders the performance trade-offs that are inherent to polyphagy untenable, and the generalist rapidly decays into an array of specialist descendants. The adaptive-specialization hypothesis doesn't rule out the possibility that specialists can speciate. But it does imply that speciation rates are much faster in generalists, and that speciation-via-specialization is the norm. On the other hand, in specialization-by-drift, generalist species stay generalist. But because a generalist species is so widespread and flexible, there is a good chance that small populations will become isolated somewhere along or beyond the edge of its geographic range. If such a population is isolated for long enough, the evolution of reproductive isolation is likely, and the evolution of host specificity may result from genetic drift and relaxed selection for polyphagy. If environments change for these specialist species, and select for broader host ranges, this selection will be inefficient, as it will be opposed by strong genetic drift, and some of the genetic underpinnings of phenotypic plasticity will have decayed. Host specificity is a dead end. Here we are contrasting one standard adaptive model with one plausible nonadaptive model; a range of other adaptive and non-adaptive hypotheses about host range are of course possible--see Discussion.

The specialization-by-drift and standard adaptive-specialization hypotheses do make some distinct and testable predictions about the phylogenetic evolution of host range. Specialization-by-drift predicts that broad host ranges will persist over evolutionary time scales; that is, host range will have a strong phylogenetic signal. By contrast, the adaptive-specialization hypothesis predicts that broad host ranges will be ephemeral, as host-use tradeoffs select against generalist phenotypes and hosts are rapidly apportioned among specialist daughter lineages. Specialization-by-drift also predicts that most cladogenetic change in host range will be asymmetrical. One of the daughter lineages produced by speciation of a polyphagous ancestor will be a host-specialist. The other will retain the host range of the ancestral population. On the other hand, standard adaptive-specialization predicts more symmetry in cladogenetic range reductions.

Comparative phylogenetic analyses of butterflies do not support the standard adaptive-specialization hypothesis: species diversification rates appear higher in specialists, host range size infrequently changes during speciation events, and when it does, expansion is as likely as contraction (Hardy and Otto 2014). Furthermore, as noted previously, polyphagy is phylogenetically conservative, at least at the level of butterfly genera (Hamm and Fordyce 2015; but see Nylin et al. 2014). Here, we estimate how host range affects species diversification in scale insects, a group for which we expect specialization-by-drift to be especially likely. Our expectation is based on three considerations: 1) Comparative phylogenetic analyses indicate a lack of evolutionary tradeoffs in host use that would select against generalist phenotypes in scale insects (Peterson et al. 2015), that is, polyphagy is cheap. 2) Scale insects have sessile adult females and disperse passively, by wind, as delicate first instar nymphs that can not survive for long away from a host. Thus, in scale insects, host specialization may be especially costly, due to high mortality during dispersal (Dixon et al. 1987). 3) Polyphagous scale insect species can have exceptionally broad host ranges, geographic ranges, and population sizes (García Morales et al. 2016; Ross et al. 2013). These are species that would seem to have an exceptionally low probability of going extinct, and a high probability of giving rise to isolated offshoots.

## Methods

Statistical tests were performed in the R software environment (R core development team 2015) on the comparative phylogenetic dataset of Hardy et al. (2015). The data are in the Dryad archive (http://dx.doi.org/10.5061/dryad.925cb). They consist of a time-calibrated, multi-locus estimate of phylogeny for 472 scale insect species, along with host range estimates for 426 of the species in the phylogeny. Hardy et al. (2015) quantified host ranges as counts of host plant families and as phylogenetic diversities (PD) of host genera.

Here, we estimated the effect of host range on taxonomic diversification rates by fitting Quantitative-state Speciation and Extinction (QuaSSE) models (FitzJohn 2010). Specifically, we compared a model in which speciation rate was a linear function of host range to a model in which speciation rate was constant and independent of host range. In both models, constant extinction rates were estimated. SSE model comparisons based on empirical phylogenies are prone to elevated Type I error rates (Madisson and FitzJohn 2014; Rabosky and Goldberg 2015). To address this issue, we used a simulation approach to adjust significance thresholds, following the recommendation of Rabosky and Goldberg (2015). Specifically, we constructed a set of 100 null datasets by simulating the evolution of a diversification-rate-neutral trait over the scale insect phylogeny, using a Brownian Motion model with the value of the σ^2^ parameter estimated from the empirical data. Then, for each simulated dataset, we calculated the difference between AIC scores (δ-AIC) of a constant-and linear-speciation-rate QuaSSE model. Lastly, we determined the significance of the empirical 5-AIC through comparison to the null distribution of simulated δ-AICs.

To assess the frequency and symmetry of cladogenetic changes in host range, we transformed the host range data to a binary trait by coding species that feed on hosts in one plant family as monophagous, and species that feed on two or more host plant families as polyphagous. Then we compared the fit of four Binary State Speciation and Extinction node-enhanced state shift (BiSSE-ness: Magnuson-Ford and Otto 2012) models: 1) a model in which there was only anagenetic evolution of host range size (this has six free parameters and is equivalent to the standard BiSSE model), 2) a model in which there is only cladogenetic evolution of host range, and the rates and symmetry of host range evolution depend on range size (this has eight free parameters), 3) a cladogenetic model in which the proportion of asymmetrical host range evolution in monophagous ancestors (i.e., the *p0a* paramter) was set to 0.5 (seven free parameters), and 4) a cladogenetic model in which in which the proportion of asymmetrical host range evolution in polyphagous ancestors (*p1a*) was set to 0.5. Full model specifications are available in the R scripts in Supplementary Materials. The significance of model comparisons was evaluated using simulation, as for the QuaSSE analyses, except with different models of neutral trait evolution. For the comparison between the anagenetic and full cladogenetic models, we used a mk2 model of discrete trait evolution to simulate neutral trait histories. For the comparisons between the full and constrained cladogenetic models, we evolved traits under an mk2 model over a version of the scale insect phylogeny in which all branch length were set to 1. Transforming branch lengths in this way approximates a history of cladogenetic (i.e., punctuated) evolution (Pagel 1994). We fit all SSE models using maximum likelihood optimization in the diversitree package (FitzJohn 2012). We also performed a Bayesian search to get an additional sense of the error of the linear-speciation-rate QuaSSE model coefficient estimates. We ran an MCMC analysis for 1000 steps, using the ML parameter estimates as starting values, and an exponential prior with rate 10. The first 100 steps were discarded as burn-in. We used Geweke's convergence diagnostic to confirm that the remaining 900 samples were from the stationary distribution (Geweke 1992; Plummer et al. 2006).

To visualize the evolution of host range over the scale insect phylogeny, we performed an ancestral state reconstruction under the estimated parameters of a BiSSE model. Lastly, we estimated the phylogenetic signal of host range using Abouheif's test (Abouheif 1999) and orthonormal decomposition (Ollier et al. 2006), implemented in the R package adephylo (Jombart et al. 2010).

## Results

We found a positive relationship between host range and diversification rate, regardless of how host range was measured, or what estimation procedure was used (Fig 1). The linear speciation model fit the data better than the constant speciation model (host family counts δ-AIC = 42.3, p-value = 0.02; PD δ-AIC = 49.7, p-value < 0.01), and had a positive slope (Table S1).

Interestingly, in the PD parameterization, the speciation rate estimates for the most host specific species were lower than the estimated extinction rate. We also found that the evolution of host range size is largely cladogenetic and strongly asymmetrical. The binary-state model with only cladogenetic evolution of host range was strongly preferred to a model with only anagenetic host range evolution (δ-AIC = 14.8, p-value < 0.01). Moreover, in the cladogenetic model, the proportion of asymmetrical host range changes in speciation events involving polyphagous ancestors was close to unity (Table S1). The full cladogenetic model was a significantly better fit to the data than a model in which cladogenetic changes in polyphagous ancestors were constrained to be symmetrical (δ-AIC = 10.5, p-value < 0.01). By constrast, constraining monophagous ancestors to be involved in only symmetrical cladogenetic changes in host range did not significantly reduce the likelihood of the model at the 0.5 level (δ-AIC = 1.7, p-value = 0.07). Note that in the best-fitting binary-state model, the full cladogenetic model, speciation rates are actually faster in specialist lineages, but so are extinction rates, and as a result, net diversification rates are an order of magnitude higher in generalists (Table S1).

We recovered strong phylogenetic signal for host range (Abouheif’s test, Dmax and SCE statistic p-values each < 0.01), although we should point out that those tests assume that host range evolves under Brownian Motion and does not affect taxonomic diversification. The phylogenetic conservatism of polyphagy is striking in the BiSSE reconstruction of ancestral host ranges (Fig 2.)

**Figure 2.**
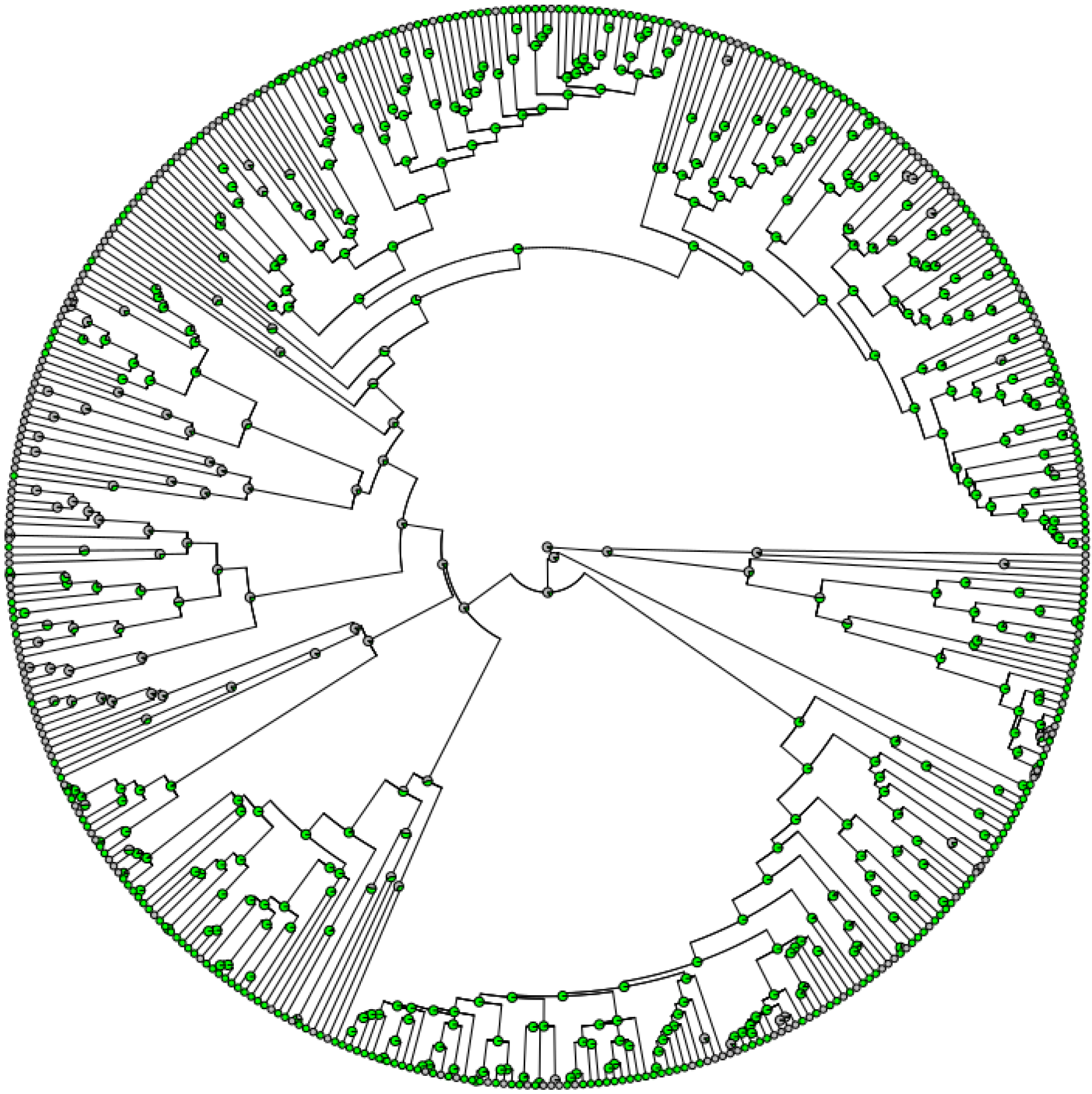
Marginal ancestral state reconstructions under the estimated BiSSE model parameters. Pie charts show proportional likelihoods at internodes. Gray: monophagy (one host plant family), Green: polyphagy (multiple host plant families).

## Discussion

In this study, we find support for two predictions of the standard adaptive-specialization hypothesis: 1) a positive relationship between host range and species diversification rate, and 2) a high proportion of cladogenetic host range reduction. This constitutes the first support for those predictions, and contrasts with the case of butterflies, which display a negative effect of host range size on diversification rate, and a low proportion of cladogenetic host range evolution (Hardy and Otto 2014). However, both of these predictions are also consistent with the specialization-by-drift hypothesis, and for scale insects, we expect specialization-by-drift to be more likely. Indeed, we recover two further patterns that are consistent with specialization-by-drift but not with the standard adaptive-specialization hypothesis: 3) asymmetry of cladogenetic host range reductions, with specialization occurring in only one of the pair of daughter species, and 4) a strong phylogenetic signal for host range, with polyphagy persisting over evolutionary time. Nevertheless, the phylogenetic patterns of host-use evolution do not tell us anything directly about the short-term adaptiveness of host specificity in scale insects, and could be explained by alternative hypotheses.

One such alternative is the serial-specialization hypothesis, first applied to parasitic tachinid flies (Stireman 2005). The starting premise is that, depending on the circumstances, selection may push for small host ranges or large ones. Examples of factors that could determine optimal host ranges for herbivorous insects include the stability of plant communities (Jaenike 1990) and the dispersal ability of the herbivores (Hardy et al. 2015). The serial-specialization hypothesis builds on that premise with two additional assumptions. First, generalists are less likely to go extinct. This is the flip side of the specialization as a dead-end hypothesis, and is supported by the fossil record and comparative phylogenetics (e.g., Jablonski 1987, Hardy and Otto 2014). Second, selection on local populations of a generalist species will often promote the evolution of host specificity and reproductive isolation. The concept of an immortal generalist species that buds off specialist daughter species is obviously something that serial-specialization shares with specialization-by-drift. And this means that phylogentic patterns of host range evolution that we estimated for scale insects are equally compatible with serial-specialization. How do we tell them apart?

Based on what else we know about scale insect biology, we think that specialization-by-drift is the more likely explanation. Specifically, in scale insects it looks like broad diets are cheap (Peterson et al. 2015), and narrow diets are expensive, owing to the mortality they are likely to cause during dispersal. But we cannot rule out the serial version of adaptive specialization; indeed, serial adaptive specialization may be highly plausible in some circumstances. The cost of host specificity in scale insects undoubtedly depends on the ecological and population genetic context (Hardy et al. 2015). For example, the costs of wind dispersal should be much lower in environments low plant species richness, or in which one plant species is extremely abundant. In fact, many specialist species of scale insects do maintain large populations on ecologically dominant hosts such as grasses and pines (García Morales et al. 2016). On the other hand, most specialist scale insect species occur in more diverse biomes and on less abundant hosts (García Morales et al. 2016). Which again points to more specialization by drift than by serial adaptation. But a glance at the phylogenetic distribution of host ranges in Figure 2 suggests that processes other than strict specialization by drift. Under the most extreme form of the specialization-by-drift hypothesis, one supreme generalist would be the progenitor of all other species, all of them specialists. In that case, barring some hard polytomies, the scale insect phylogeny would be completely unbalanced, and the generalist would be sister to its most recent offshoot (Stireman 2005). That's not what Figure 2 looks like.

For other insects, especially insects with more efficient dispersal modes, host specialization by drift may be less likely. For example, in butterflies, the vagility, longevity and complex sensory systems of adult females may dramatically lower costs for host-plant specialization, and could open the door to more adaptive specificity. On the other hand, the costs of polyphagy are as unclear for butterflies as they are for scale insects, and that leaves the door open for non-adaptive specificity. Butterfly life history may also serve to make specialization-by-drift harder to detect; it may be more likely that specialization by drift is followed by adaptive niche transformation and expansion, a process that would be consistent with the symmetrical evolutionary changes in host range estimated by Hardy and Otto (2014). But the most likely candidates for specialization-by-drift are other groups for which specificity is especially costly; that is, groups with low-mobility adult females and haphazard dispersal such as root-feeding weevils (Curculionidae: Entiminae), stick insects (Phasmatodea), and bagworm moths (Psychidae) (Normark and Johnson 2011).

In our description of specialization-by-drift we assumed an allopatric mode for speciation. Alternative non-adaptive modes of speciation, such as conflictual speciation (Crespi and Nosil, 2013), might also be relevant. This is especially true for scale insects, which engage in obligate endosymbioses that are expected to increase the potential for genetic conflict (Brucker and Bordenstein 2012). However, unless strong barriers to hybridization are initially present, genes involved in conflicts between scale insects and their endosymbionts might actually homogenize gene pools, and forestall speciation (Crespi and Nosil 2013). Which mean that genetic conflicts probably play a more important role in crystallizing the genetic divergences which occurred in allopatry, than in driving sympatric speciation.

Our results imply that, for most scale insects, ecological specificity is a dead-end. The evidence is mixed for ecological specificity being a dead-end in other lineages (Jablonski 2008; Day et al. 2016). This is not surprising, given the dissonance in the theory for how specificity should affect macroevolution. On the one hand, we have the dead-end. On the other, the master of a trade. As Jablonski (2008) points out in his excellent review of species selection, most of the traits that we expect to increase the risk of extinction should also increase the chance of speciation. Ecological specificity falls into this category. Furthermore, if specialization increases competitive ability, it might actually decrease the risk of extinction (Roberts and Hawkins 1999; Stireman 2005). The net effect that it has on species diversification should depend on the relative costs of big niches (e.g., inefficient metabolism) and small niches (e.g., smaller, patchier resources), which should vary among species, depending on factors such as dispersal mode and environmental heterogeneity.

Our results are contingent on the parameters that were included in the models of trait-dependent diversification and on the assumptions made by those models. Host range *per se* cannot be the most proximate factor affecting speciation and extinction dynamics. Rather, host range should affect diversification by influencing population genetic parameters like population size and structure, by playing a role in structuring the selective environment, and by affecting the probability of colonizing new hosts and new places (Jahner et al. 2011; Slove and Janz 2011). In this study we fail to reject the hypothesis that host specialization is caused by genetic drift in scale insects. Further research is needed to test other aspects of that hypothesis, for example comparative assessments of population size and structure in generalist and specialist species, and experiments designed to assess the extent of drift, or the strength of selection at loci important for host use and across the genome.

## Conclusions

Most herbivorous insect species are host-plant specialists. The idea that a specialist on its own host should be able to outperform generalists is intuitively appealing, but it fails to account for how host specificity may alter population genetic factors and the efficiency of natural selection. It also fails to align with the ever-expanding body of research that indicates the superior performance of generalists, and fails to predict the distribution of host range across herbivorous insect phylogenies (e.g., Hardy & Otto 2014, Peterson et al. 2015). Here we find patterns of host range evolution that are consistent with a model of specialization by drift. Population genetic processes other than natural selection should be more fully and more routinely incorporated into ideas about the species diversification of the most species-rich metazoans.

## Acknowledgments

Our collaboration was fostered by travel funds provided by NSF (DEB-1258001 and EF-1115191). Further support for this research was provided by the Alabama Agricultural Experiment Station to NBH. This manuscript was greatly improved by the insightful comments of the editors and anonymous reviewers. We are grateful for their help.

## References

Abouheif, E. 1999. A method for testing the assumption of phylogenetic independence in comparative data. Evol. Ecol. Res. 1:895–909.

Agosta, S. J., and J. A. Klemens. 2009. Resource specialization in a phytophagous insect: no evidence for genetically based performance trade-offs across hosts in the field or laboratory. J. Evol. Biol. 22:907–912.

Brucker, R.M. and S. R. Bordenstein. 2012. Speciation by symbiosis. Trends Ecol. Evol. 27: 443–451.

Cope, E. D. 1896. The primary factors of organic evolution. Open Court Publishing CO., Chicago.

Crespi, B. and P. Nosil. 2013. Conflictual speciation: species formation via genomic conflict. Trends Ecol. Evol. 28: 48–57.

Day, E. H., X. Hua and L. Bromham. 2016. Is specialization an evolutionary dead end? Testing for differences in speciation, extinction and trait transition rates across diverse phylogenies of specialista and generalists. J. Evol. Bio. DOI: 10.1111/jeb.12867

Dixon, A. F. G., P. Kindlmann, J. Lepš, and J. Holman. 1987. Why there are so few species of aphids, especially in the tropics. Am. Nat. 129:580–592.

Ehrlich, P. R., and P. H. Raven. 1964. Butterflies and plants: a study in coevolution. Evolution 586–608.

FitzJohn, R. G. 2012. Diversitree: comparative phylogenetic analyses of diversification in R. Methods Ecol. Evol. 3:1084–1092.

FitzJohn, R. G. 2010. Quantitative traits and diversification. Syst. Biol. 59:619–633.

Forister, M. L., A. G. Ehmer and D. J. Futuyma. 2007. The genetic architecture of a niche: variation and covariation in host use traits in the Colorado potato beetle. J. Evol. Bio. 20:985–996.

Forister, M. L., V. Novotny., A. K. Panorska, L. Baje, Y. Basset, P. T. Butterill, L. Cizek, P. D. Coley, F. Dem, I. R. Diniz and P. Drozd. 2015. The global distribution of diet breadth in insect herbivores. Proc. Natl. Acad. Sci. 112:442–447.

Futuyma, D. J., and A. A. Agrawal. 2009. Macroevolution and the biological diversity of plants and herbivores. Proc. Natl. Acad. Sci. 106:18054–18061.

Futuyma, D. J., and G. Moreno. 1988. The evolution of ecological specialization. Annu. Rev. Ecol. Syst. 19:207–233.

García Morales, M., B. D. Denno, D. R. Miller, G. L. Miller, Y. Ben-Dov and N. B. Hardy. 2016. ScaleNet: a literature-based model of scale insect biology and systematics. Database 2016:bav118, doi: 10.1093/database/bav118

Geweke, J. Evaluating the accuracy of sampling-based approaches to calculating posterior moments. In Bayesian Statistics 4 (ed JM Bernado, JO Berger, AP Dawid and AFM Smith). Clarendon Press, Oxford, UK.

Gompert, Z., J. P. Jahner, C. F. Scholl, J. S. Wilson, L. K. Lucas, V. Soria-Carrasco, J. A. Fordyce, C. C. Nice, C. A. Buerkle, and M. L. Forister. 2015. The evolution of novel host use is unlikely to be constrained by tradeoffs or a lack of genetic variation. Mol. Ecol. 24:2777–2793.

Hamm, C. A., and J. A. Fordyce. 2015. Patterns of host plant utilization and diversification in the brush-footed butterflies. Evolution 69:589–601.

Hardy, N. B. and L. G. Cook. 2010. Gall-induction in insects: evolutionary dead-end or speciation driver? BMC Evol. Bio. 10:257; DOI: 10.1186/1471-2148-10-257

Hardy, N. B., and S. P. Otto. 2014. Specialization and generalization in the diversification of phytophagous insects: tests of the musical chairs and oscillation hypotheses. Proc. R. Soc. B Biol. Sci. 281:20132960.

Hardy, N. B., D. A. Peterson and B. B. Normark. 2015. Scale insect host ranges are broader in the tropics. Bio. Let. 11: 20150924; DOI: 10.1098/rsbl.2015.0924

Huxley, J. 1942. Evolution: the modern synthesis. George Allen and Unwin, London.

Jaenike, J. 1990. Host specialization in phytophagous insects. An. Rev. Ecol. Sys. 21: 243–273.

Jablonski, D. 1986. Background and mass extinctions: The alteration of macroevolutionary regimes. Science 231: 129–133.

Jablosnki, D. 2008. Species selection: Theory and Data. Annu. Rev. Ecol. Evol. Syst. 39: 501–524.

Jahner, J. P., M. M. Bonilla, K. J. Badik, A. M. Shapiro and M. L. Forister 2011. Use of exotic hosts by Lepidoptera: widespread species colonize more novel hosts. Evolution 65: 2719–2724.

Janz, N., S. Nylin, and N. Wahlberg. 2006. Diversity begets diversity: host expansions and the diversification of plant-feeding insects. BMC Evol. Biol. 6:4.

Jombart, T., F. Balloux, and S. Dray. 2010. adephylo: new tools for investigating the phylogenetic signal in biological traits. Bioinformatics 26:1907–1909.

Kelley S. T., and B. D. Farrell. 1998. Is specialization a dead end? The phylogeny of host use in Dendroctonus bark beetles (Scolytidae). Evolution 52:1731–1743.

Maddison, W. P., and R. G. FitzJohn. 2015. The Unsolved Challenge to Phylogenetic Correlation Tests for Categorical Characters. Syst. Biol. 64:127–136.

Magnuson-Ford, K., and S. P. Otto. 2012. Linking the investigations of character evolution and species diversification. Am. Nat. 180:225–245.

Mitter, C., B. Farrell, and B. Wiegmann. 1988. The Phylogenetic Study of Adaptive Zones: Has Phytophagy Promoted Insect Diversification? Am. Nat. 132:107–128.

Normark, B. B., and N. A. Johnson. 2011. Niche explosion. Genetica 139:551–564.

Nylin, S., J. Slove and N. Janz. 2014. Host plant utilization, host range oscillations and diversification in nymphalid butterflies: a phylogenetic investigation. Evolution 68:105–124.

Ollier, S., P. Couteron, and D. Chessel. 2006. Orthonormal Transform to Decompose the Variance of a Life-History Trait across a Phylogenetic Tree. Biometrics 62:471–477.

Pagel, M. 1994. Detecting correlated evolution on phylogenies: a general method for the comparative analysis of discrete characters. Proc. R. Soc. Lond. B. 255:37–45.

Peterson, D. A., N. B. Hardy, G. E. Morse, I. C. Stocks, A. Okusu, and B. B. Normark. 2015. Phylogenetic analysis reveals positive correlations between adaptations to diverse hosts in a group of pathogen-like herbivores. Evolution 69:2785–2792.

Plummer, M., N. Best, K. Cowles and K. Vines. 2006. CODA: Convergence diagnosis and output analysis for MCMC. R news 6:7–11.

Rabosky, D. L., and E. E. Goldberg. 2015. Model Inadequacy and Mistaken Inferences of Trait-Dependent Speciation. Syst. Biol. Syu131.

Rainford, J.L. and Mayhew, P.J., 2015. Diet evolution and clade richness in Hexapoda: a phylogenetic study of higher taxa. Am. Nat. 186:777–791.

Roberts, C. M. and J. P. Hawkins. 1999. Extinction risk in the sea. Trends Ecol. Evol. 14: 241–246.

Simpson, G. 1953. The major features of evolution. Colombia Univ. Press, New York.

Slove, J., and N. Janz. 2011. The relationship between diet breadth and geographic range size in the butterfly subfamily Nymphalinae—a study of global scale. PLoS ONE 6:e16057.

Stireman, J. O. 2005. The evolution of generalization? Parasitoid flies and the perils of inferring host range evolution from phylogenies. J. Evol. Bio. 18: 325–336.

Wiens, J. J., Lapoint, R. T. and Whiteman, N.K. 2015. Herbivory increases diversification across insect clades. Nature communications 6:8370, DOI: 10.1038/ncomms9370.

Yoder, J.B., E. Clancey, S. Des Roches, J. M. Eastman, L. Gentry, W. Godsoe, T. J. Hagey, D. Jochimsen, B. P. Oswald, J. Robertson and B. A. J. Sarver. 2010. Ecological opportunity and the origin of adaptive radiations. J. Evol. Bio. 23: 1581–1596.

